# *Escherichia coli* extra-intestinal population translocation in leukemia patients

**DOI:** 10.1101/2024.01.26.577459

**Authors:** Julie Marin, Violaine Walewski, Samira Dziri, Mélanie Magnan, Erick Denamur, Etienne Carbonnelle, Antoine Bridier-Nahmias

## Abstract

*Escherichia coli*, a commensal species of the human gut, is an opportunistic pathogen which can reach extra-intestinal compartments, including the bloodstream and the bladder, among others. In non-immunosuppressed patients, purifying or neutral evolution of *E. coli* populations has been reported in the gut. Conversely, it has been suggested that when migrating to extra-intestinal compartments, *E. coli* genomes undergo diversifying selection as supported by strong evidence for adaptation. The level of genomic polymorphism and the size of the populations translocating from the gut to extra-intestinal compartments is largely unknown.

To gain insights in the pathophysiology of these translocations, we investigated the level of polymorphism and the evolutionary forces acting on the genomes of 77 *E. coli* isolated from various compartments in three immunosuppressed patients. We detected a unique strain for each patient across the blood, the urine and the gut. In one case, all isolates recovered were mutators i.e. isolates with a very high mutation rate. In all instances, we observed that translocation encompasses the majority of the genomic diversity present in the gut. The same signature of selection, whether purifying or diversifying, and as anticipated, neutral for mutator isolates, was observed in both the gut and bloodstream. Additionally, we found a limited number of non-specific mutations among compartments for non-mutator isolates. In all cases, urine isolates were dominated by neutral selection. These findings indicate that substantial proportions of populations are undergoing translocation and that they present a complex compartment-specific pattern of selection at the patient level.

**Importance:** It has been suggested that intra and extra-intestinal compartments differentially constrain the evolution of *E. coli* strains. Whether host particular conditions, such as immunosuppression, could affect the strain evolutionary trajectories remain understudied. We found that, in immunosuppressed patients, large fractions of *E. coli* gut populations are translocating with variable modifications of the signature of selection for commensal and pathogenic isolates according to the compartment and/or the patient. Such multiple site sampling should be performed in large cohorts of patients to get a better understanding of *E. coli* extra-intestinal diseases.

## Introduction

Bloodstream infections (BSIs) are still a major concern among onco-hematologic patients, depending on the type of pathogen, level of host immunodeficiency and status of underlying disease. Despite advancements in the clinical management of hematological malignancies, BSIs remain life-threatening complications in the clinical course of these patients, with reported crude mortality rate up to 40% (1–3). A clear shift of bacterial species causing BSIs in patients with hematological malignancies has been recently reported, transitioning from Gram-positives to Gram-negatives. In the first place Enterobacteriaceae, and in particular *Escherichia coli*, represent the most frequently involved bacterial species, together with the worrisome and growing phenomenon of multiresistant bacteria (4, 5).

*E. coli* is a commensal species of the lower intestine of humans (6). The gut is the primary habitat of *E. coli*, and probably the main ecological context of selection. Virulence genes, for instance, are thought to be primarily selected in the intestine as a by-product of commensalism (7, 8). *E. coli* is also an opportunistic pathogen, frequently responsible for intestinal and extra-intestinal infections (9). When reaching a new compartment, such as the bladder or the bloodstream, *E. coli* faces new challenges and new opportunities for adaptation.

In the past, only a few studies have investigated the signatures of selection in commensal *E. coli* isolates and in isolates sampled from extra-intestinal infections, revealing different scenarios. The evolution of commensal *E. coli* has been found to be governed by purifying selection, whether it implies the entire genome (10) or specific genes such as the H7 flagellin genes (11). However, another study, following the evolution of a clone in a single individual over almost a year, did not find any evidence for selection in the gut (12). On the contrary, it was found that adaptation is at play during chronic and acute infections (13). In particular, it was shown that *E. coli* isolates colonizing extra-intestinal sites were adapting under strong selective pressure, with an excess of non-synonymous mutations and patterns of convergence at the gene level (including for H7 flagellin genes). Evidence for adaptation during chronic infection have been emphasized for other bacteria, such as *Burkholderia dolosa* (14) or *Pseudomonas aeruginosa* (15–17), with genotypically and phenotypically diversifying lineages during cystic fibrosis infections for instance (18–21). Adaptation to the human host (22, 23), evasion from the immune response (24, 25) and acquisition of antibiotic resistances (26, 27) are favored by this accumulation of mutations. In addition, niche adaptation shapes the allelic diversity among compartments with specific mutations occurring in the gut or in the bladder associated with functions increasing *E. coli* fitness in one or the other compartment (28).

What happens when the intestinal barrier and the immune response are weakened? Patient’s deficiencies, such as immunosuppression, weakening of the intestinal barrier by antibiotic therapy or chemotherapy, can increase the risk of *E. coli* extra-intestinal infection (29–31). Anticancer chemotherapy drugs have a direct impact on the intestinal microbiota (32). The resulting dysbiosis can lead to intestinal mucositis which increases the risk of bacterial translocation to the bloodstream (33). Moreover, among patients undergoing chemotherapy, leukemia patients are highly prone to extra-intestinal infections and relapse (34). The weakening of intestinal barriers and the lack of immune defense could modify the adaptive conditions described for commensal and pathogenic isolates of *E. coli* in non-immunosuppressed patients. Modifications in the signature of selection are therefore expected for commensal and pathogenic isolates of *E. coli* in immunosuppressed patients. In addition, the level of genomic polymorphism and the proportion of the population translocating is largely unknown.

Here, we evaluated the selective forces acting on *E. coli* evolution in three immunosuppressed patients among three compartments, the bladder, the bloodstream and the gut. For each patient, we analyzed whole genome sequences of the isolates found in each compartment to evaluate genetic diversity and assess the strength of the various selective processes at play.

## Material and methods

### Sampling

Clinical isolates were isolated from three patients with hematologic disease and *E. coli* sepsis hospitalized in Avicenne (Seine-Saint-Denis, France). Patient A had three infectious episodes while patient B and C had one each. For each episode, isolates were sampled from positive blood culture along from bladder and feces samples on the same or subsequent day. The initial blood culture was obtained before any antibiotic therapy. All procedures performed were in accordance with the ethical standards of the responsible committee on human experimentation (institutional and national), validated by the ethics committee of Avicenne’s hospital (Comité Local d’Ethique d’Avicenne). Information on the type and status of hematologic disease, presence of neutropenia (<0.5×10^9^/L), previous exposure to any antibiotic therapy, including prophylaxis or treatment of prior infectious episodes, type of infection, microbiological isolate, and outcome was collected in a database.

### Patient characteristics

The three patients (A to C) studied had hematological pathologies: two acute myeloid leukemia and one with Hodgkin’s lymphoma. Patient A was diagnosed with acute myeloid leukemia and myelofibrosis in January 2014 (Acute myeloblastic leukemia with minimal maturation M1). He was treated with allograft and experienced relapse in March 2015, with febrile neutropenia and presence of circulating blasts. During the BSI, the patient presented an inflammatory syndrome, a severe immunosuppression and had received antibiotics in the previous 30 days. Patient B also suffered from undifferentiated acute myeloblastic leukemia (M0) diagnosed and allografted in 2015. At the time of BSI, he was receiving chemotherapy, had severe immunosuppression, an inflammatory syndrome, and had received antibiotics within the preceding 30 days. Patient C was diagnosed with Hodgkin’s lymphoma in 2009. Relapsed in 2017 during the BSI episode, the patient exhibited inflammatory syndrome, severe immunosuppression and had previously received antibiotics.

### Prior in vitro phylogroup typing

We determined the phylogroup of each sample (blood, urinary or feces) using quadruplex PCR (35) to select isolates belonging to the same phylogroup for each infection episode.

### Genome sequencing and assembling

We whole-genome sequenced each sample with Illumina technology (MiSeq and HiSeq 2500) using Nextera XT library preparation kits as instructed by the manufacturer (Illumina, San Diego, CA). Fastq files (raw sequencing data) were submitted to the European nucleotide archive (see **Table S1** for accession numbers). Genome assembly was performed with SPAdes v.3.15.5 (36) .

For each patient, one strain was chosen as a reference and was sequenced with the Oxford Nanopore Technologies MinION platform using an R9.4 flow cell. We prepared the samples with kits LSK-108 and NBD-104 for library preparation and barcoding. The Nanopore reads were filtered with Filtlong (37) using the following parameters: minimum length of 1000 and 95% of the best reads kept. High-quality assemblies of the three reference genomes were obtained through a hybrid strategy using both Illumina and Nanopore reads with Unicycler V0.4.4 (38).

### Core and accessory genome

The 77 SPAdes assemblies were annotated with Prokka (39). Plasmid sequences were predicted by PlaScope (40). We then performed pan-genome analysis from annotated assemblies with Roary on the 77 genomes using default parameters (41). The core genome alignment and the list of genes of the accessory genome were generated for the three patients.

### Variant calling (SNPs and deletions)

We first discarded bases with a low-quality score (< 30) and removed the adapters with Trim Galore [a wrapper of the Cutadapt program (42)]. To detect SNPs, we aligned the reads of each patient to the corresponding reference sequence (**Table S1**) with Snippy 4.4.0 (43) with the following parameters: the nucleotide minimum quality to be analyzed (basequal) equal to 20, the minimum number of reads covering a site (mincov) equal to 10 and the minimum proportion of those reads different from the reference (minfrac) equal to 0.9. To detect structural variants (deletions) we first mapped the reads to the corresponding reference assembly with BWA-MEM (44) and then analyzed the obtained Sequence Alignment Map (sam) files with Whamg (45). As recommended, we remove calls smaller than 50 bp and larger than 2 Mbs. For each strain, we also removed calls with less than 5 supporting reads.

### Typing and genotypic antibiotic resistance and virulence

We used an in-house script, petanc (46), that integrates several existing bacterial genomic tools to perform the typing of isolates with several genotyping schemes using the genomic tool SRST2 (47). Sequence types (STs) were defined using the Warwick MLST scheme and the Pasteur scheme (48, 49). We only used the Warwick scheme for the analyses described thereafter. We also determined the O:H serotypes and the FimH alleles (Ingle et al. 2016; Roer et al. 2017). The phylogroups were confirmed using the ClermonTyping method (50).

The resistome and virulome were first established using the in-house script Petanc (46). They were defined by BlastN with Abricate (https://github.com/tseemann/abricate) using the ResFinder (version 4.2.2) database (51), a custom database including the VirulenceFinder database and VFDB (52, 53), to which we added selected genes (54). We set the threshold for minimum identity to 80% with a minimum coverage of 90%. Next, we built a pan-resistome and pan-virulome including all the antibiotic resistance and virulence associated genes of all isolates. We mapped the reads to the corresponding pan-resistome and pan-virulome with BWA-MEM (44). We considered a gene as present when we found more than 80% coverage and at least one read. We also tested more conservative thresholds with 80% coverage and more than 5 reads.

### Genomic diversity and traces of selection

We compared the rates of nonsynonymous and synonymous mutation by computing dN/dS ratios (R language (55)) from the gene alignments obtained with Roary, to evaluate the genomic traces of selection. First, we removed the identical sequences. Next, we calculated the number of non-synonymous substitutions (dN) and synonymous substitutions (dS) as the observed number of substitutions of each type divided by the number of potential substitutions of the same type in the considered sequence. For each codon, the number of potential non-synonymous or synonymous substitutions is determined by the genetic code. We then computed the dN/dS ratio for each pair of sequences, we determined the median dN/dS ratio and assess whether it significantly differed from 1 (Wilcoxon test), where 1 represents perfect balance between diversifying and purifying selection indicating no visible selection. When in a pair of sequences 1 or more non synonymous and 0 synonymous mutations were encountered, an infinite value is returned. Those were replaced by the maximum non-infinite value obtained for the considered set of sequences. ‘Not a number’ (NaN) values, when there was no non-synonymous mutation, were replaced by 0.

## Results

We evaluated the genomic diversity (SNPs and deletions) and the genomic traces of selection in 77 *E. coli* isolates sampled concomitantly in the gut, blood and urine compartments of three patients (**Figure 1**). The distribution of isolates for each patient and compartment is detailed in **Table 1**.

**Figure 1.**
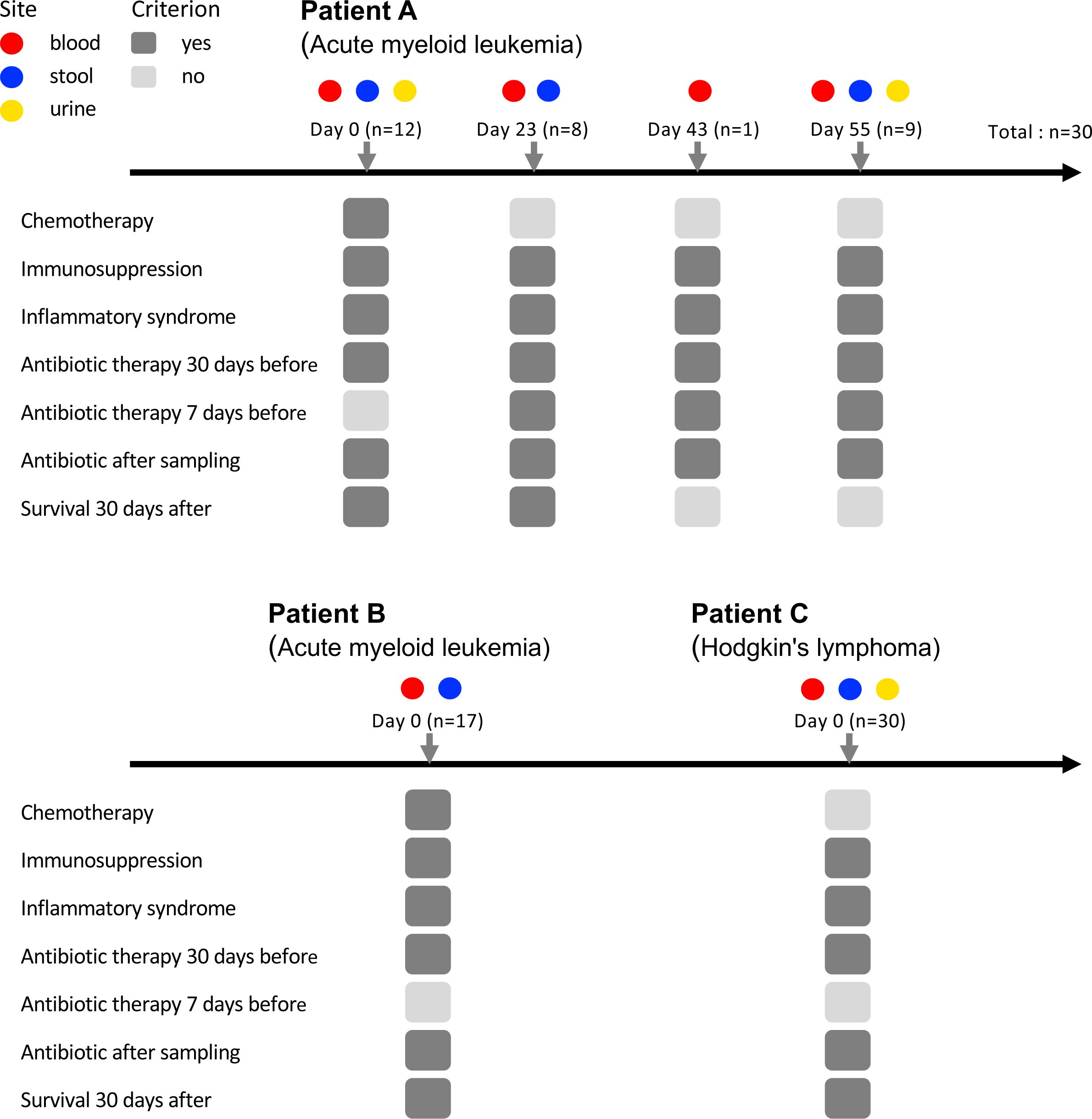
Patient follow-up with their main clinical characteristics and the sampling schemes. Absence (light gray) or presence (dark grey) of each criterion are shown. Patient A had four infectious episodes, patients B and C had one infectious episode.

**Table 1.** Patient sampling and strain typing.

| Patient | Number of isolates<br>(stool – blood – urine) † | Phylogroup | Sequence type | Serotype | *fimH* allele |
| --- | --- | --- | --- | --- | --- |
| Patient A | 30 (13 -12 - 5) | A | 10 | O101:H9 | *fimH54* |
| Patient B* | 17 (10 - 7) | B1 | 1431 | O8:H19 | *fimH32* |
| Patient C | 30 (10 - 10 - 10) | B2 | 131 | O25b:H4 | *fimH30* |

Phylogroups, MLSTs (Warwick and Pasteur), serotypes and *fimH* alleles are indicated. All the isolates of patient B were mutators (\*).
(†) Patients B and C were sampled at a single time point whereas samples of patient A correspond to 4 timepoints.

### Phylogroup, ST, serotype and fimH allele diversity

Within each infection episode, all isolates belonged to the same phylogroup and ST and had the same serotype and *fimH* allele (**Table 1**). Isolates belonged to the phylogroup A (patient A), B1 (patients B) and B2 (patient C).

### Global genomic diversity

We first looked at the SNP diversity. At the exception of patient B, a low number of mutated genes (between 8 and 11) and no deletion was recovered when comparing isolates of each patient between them (**Table S2**). For patient B, we found 328 mutated genes. Indeed, for patient B isolates we noted a deletion of *mutS* (see below) and non-synonymous mutations in *mutL*). These inactivations of the DNA mismatch repair system conferred to them a mutator phenotype (56) (**Table S3**).

In line with those results, we found a low number of SNPs for patients A and C (mean number of 5.07 [95% confidence interval: 4.20-5.95]) compared to patient B (mean number of 117.13 [82.16-152.10], p-value = 1.237e-08 (Wilcoxon test)) (**Figure 2**). Similarly, we did not find any difference between intra- and extra-intestinal isolates among variant categories (**Table 2**) (chi-squared test after correcting for sample size, p-value = 0.99). No common gene with SNPs was identified among patients (**Table S2 and S4**).

**Figure 2.**
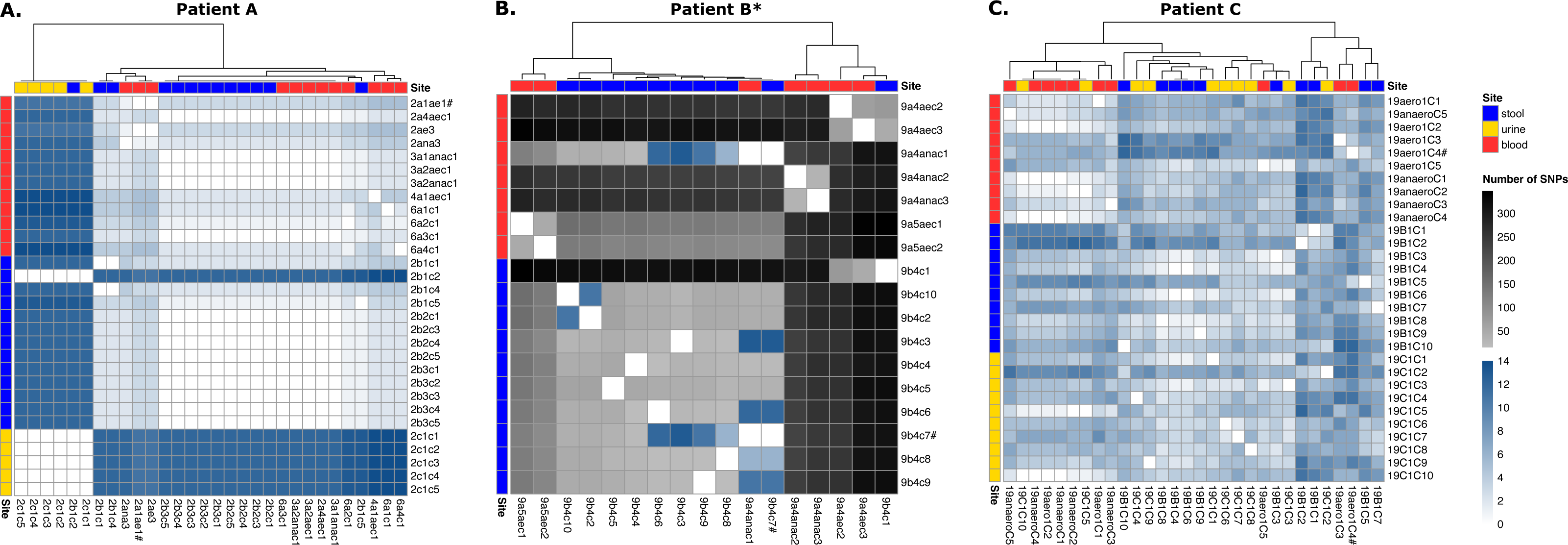
Genomic diversity of *E. coli* isolates among samples and compartments for the three patients. (A-C) Heatmaps showing the number of SNPs between each isolate of patients (A, B and C). Samples are ordered by site horizontally and clustered according to their SNP similarities vertically (method: complete). All isolates of patient B are mutators (*). The isolate corresponding to the reference sequence is indicated (#). We used the same color scale for all patients. Note that the number of SNPs for patients A and C is between 0 and 14 and between 0 and 400 for patient B.

**Table 2.** Number of structural variant types among isolates of a patient’s infection episode found with snippy.

|  | Site | conservative<br>inframe<br>insertion | disruptive<br>inframe<br>deletion | frameshift<br>variant | missense<br>variant | non coding<br>transcript<br>variant | stop<br>gained | stop lost | synonymous<br>variant |
| --- | --- | --- | --- | --- | --- | --- | --- | --- | --- |
| **Patient A** | blood<br>(n=12) | 9 | 0 | 1 | 3 | 0 | 1 | 0 | 9 |
|  | stool<br>(n = 13) | 12 | 1 | 0 | 6 | 0 | 0 | 1 | 17 |
|  | urine<br>(n=5) | 0 | 5 | 0 | 20 | 0 | 0 | 5 | 20 |
| **Patient B\*** | blood<br>(n=7) | 0 | 0 | 216 | 541 | 5 | 11 | 20 | 292 |
|  | stool<br>(n=10) | 0 | 0 | 86 | 177 | 1 | 4 | 4 | 146 |
| **Patient C** | blood<br>(n=10) | 0 | 0 | 9 | 7 | 0 | 0 | 0 | 2 |
|  | stool<br>(n=10) | 0 | 0 | 16 | 26 | 0 | 0 | 0 | 14 |
|  | urine<br>(n=10) | 0 | 0 | 12 | 20 | 0 | 0 | 0 | 6 |
| **Intra-intestinal**<br>**isolates** | stool<br>(n=23) | 12 | 1 | 16 | 32 | 0 | 0 | 1 | 31 |
| **Extra-intestinal**<br>**isolates** | blood/urine<br>(n=37) | 9 | 5 | 22 | 50 | 0 | 1 | 5 | 37 |

We excluded patient B from intra- and extra-intestinal isolates because all the 17 isolates were mutators (\*). n: number of isolates.

### Within patient compartment diversity

All isolates from a given patient were highly similar (except for patient B with mutator isolates) (**Figures 2-3**). As a consequence, no large deletions were detected because we used one of these isolates as the reference strain for each patient (**Table S2**).

**Figure 3.**
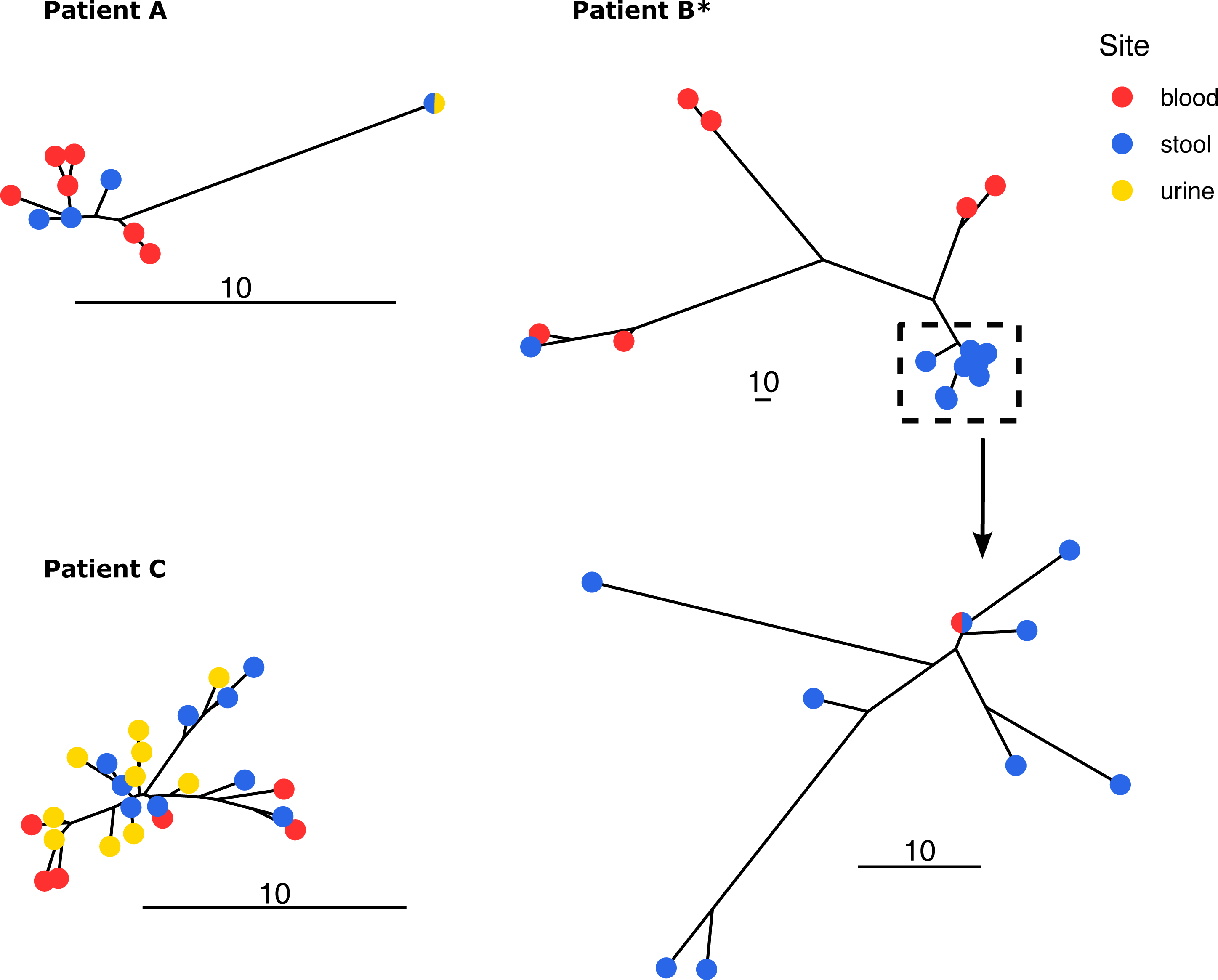
Unrooted trees of *E. coli* isolates from patients A, B and C. Trees were built using neighbor joining from the substitution presence/absence matrix. The scale indicates the number of substitutions. All isolates of patient B are mutators (*). We zoomed in on a clade of the patient B to highlight the scale difference. The bicolor points (patients A and C) denote the presence of isolates sampled in different sites with identical sequences (0 SNP).

The majority of SNPs were shared among compartments (**Figure 4**). For patient A, 13 over 20 SNPs were identical among at least 2 compartments and 13 over 22 for patient C. For patient B (mutator strain), despite a high level of polymorphism, most of the SNPs were shared among stool and blood isolates. Furthermore, most of the stool polymorphism was found in other compartments, 87%, 77% and 87% for patient A, B and C respectively, suggesting a large population translocation from the gut.

**Figure 4.**
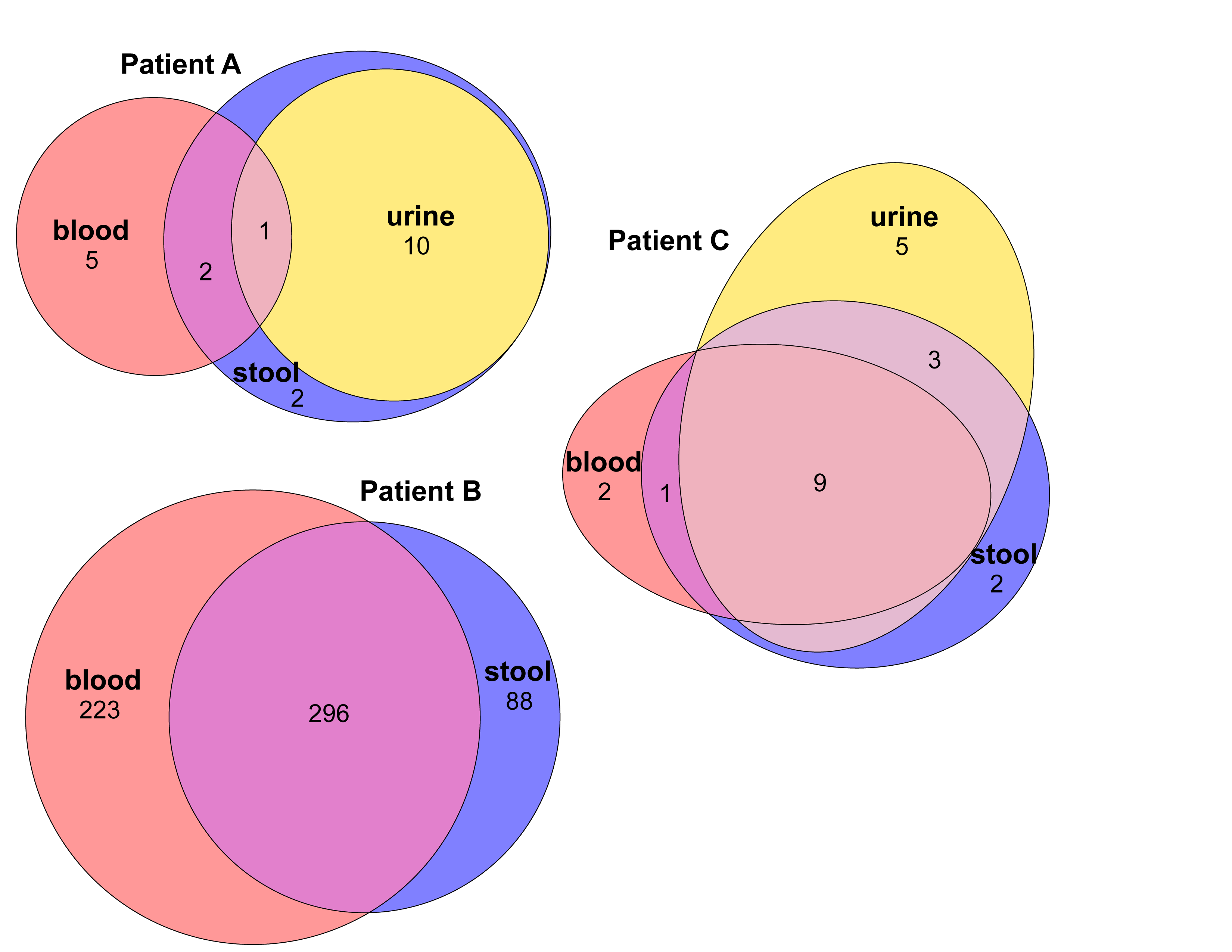
Venn diagrams showing the SNPs distribution among compartments for patients A, B and C. The ellipses are proportional to the number of SNPs.

### Antibiotic resistance and virulence associated gene content

Regarding genes associated with antibiotic resistance, isolates within each infection were highly similar. We only found few differences among isolates of patient B (mutator isolates) and A which were sampled at different times (**Figures 1 and 5**). For patient A, we found gene presence/absence discrepancies for three resistance genes, *sul2*, *tet(A)* and *dfrA5*. The gene *sul2* was consistently predicted as plasmidic across all isolates (**Table S5**). The genes *dfrA5* and *tet(A)* were found on the same contig in 20 out of 23 isolates possessing both genes, and were predicted as plasmidic for 96.67% and 86.96% of the isolates respectively. For patient B, we found presence/absence discrepancies for all the 7 resistance genes detected. Specifically, the genes *qnrS1*, *bla*CTX-M-15, *bla*TEM-1B, *aph*(6)-Id, *aph*(3’’)-Ib and *sul2* were always found on the same contig and predicted as plasmidic. The gene *dfrA*14 was predicted as plasmidic for all isolates. Similar results were obtained using a more conservative threshold (at least 5 reads covering more than 80% of the gene) (**Figure S1**), with the exception of the gene *mdf(A)*, predicted as chromosomic, which was missing for three isolates of patient B.

**Figure 5.**
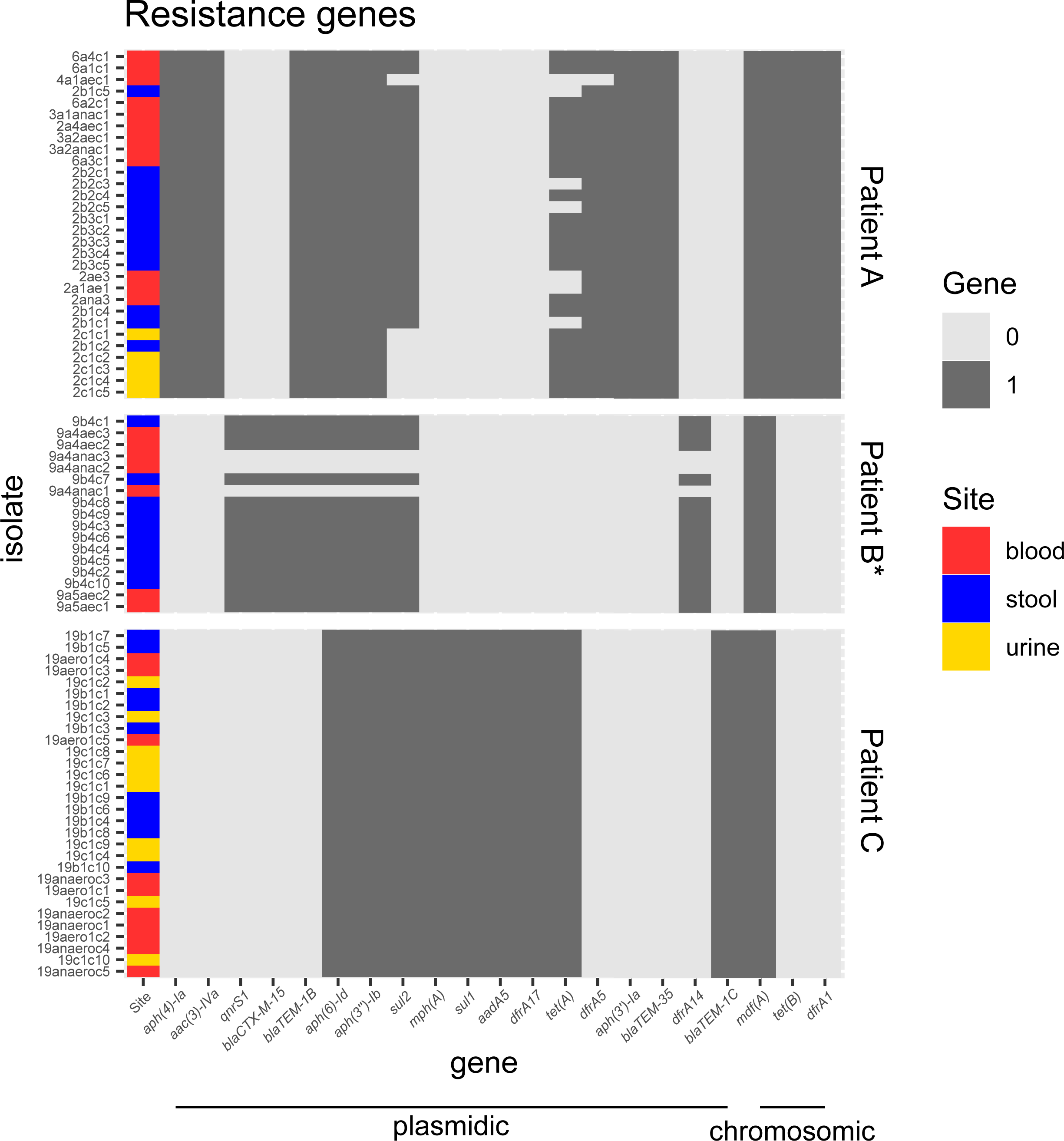
Presence/absence heatmaps of antibiotic resistance genes of *E. coli* isolates when compared to the pan-resistome (including the resistance genes of all isolates). We considered a gene as present when at least 80% of its length was covered by at least one read. Genes are ordered by synteny on contigs. All isolates of patient B are mutators (*). The prevailing predicted localization of genes by PlaScope (chromosomic or plasmidic) is indicated (full list in supplementary Table S5). Note that chromosomal genes are not mobile.

We also found high gene content similarity for virulence associated genes (**Figure 6**). An identical gene content was found in patient A isolates. For patient C, there was a discrepancy for a single gene, *iss*11, consistently predicted as chromosomic, and absent in four isolates. Slightly more differences were found for patient B (mutator strain). The genes *espX5*, *espX1* and *gad*20 were missing for 1, 1 and 5 isolates respectively, all predicted as chromosomic (**Table S6**). We found more discrepancies with a more conservative threshold (at least 5 reads covering more than 80% of the gene), highlighting the lower sequencing depth of two isolates in particular (**Figure S2**). In patient A, all the isolates had the same gene content as the exception of 2 isolates (2ana3 and 9b4c8). We found 6 missing genes for 2ana3 and one missing gene for 2c1c5. These genes were all predicted as chromosomic. In addition, *entA*, *entB* genes were always adjacent on the same contig (**Table S6**). In patient C, the gene *iss*11, always predicted as chromosomic, was missing for 5 isolates. We found more differences for patient B. Over the 47 virulence-associated genes detected, all predicted as chromosomic, only 15 were present in all isolates.

**Figure 6.**
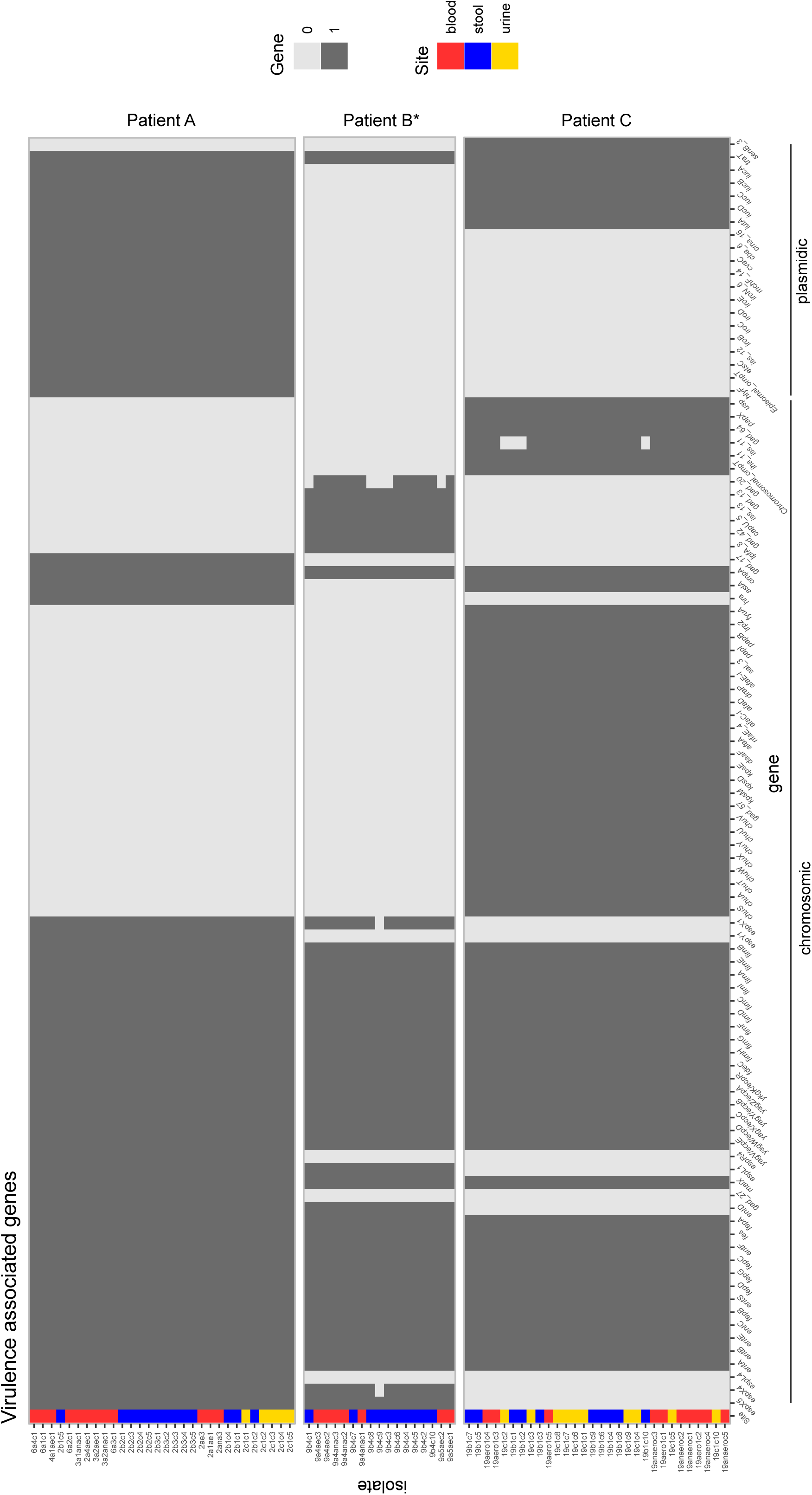
Presence/absence heatmaps of virulence associated genes of *E. coli* isolates when compared to the pan-virulome (including the virulence genes of all isolates). We considered a gene as present when at least 80% of its length was covered by at least one read. Genes are ordered by synteny on contigs. All isolates of patient B are mutators (*). The prevailing predicted localization of genes by PlaScope (chromosomic or plasmidic) is indicated (full list in supplementary Table S6). Note that plasmidic genes are not mobile, at the opposite of resistance genes (see Figure 5).

### Genomic traces of selection

We computed dN/dS ratios to test whether gene sequences evolved neutrally or were under purifying or diversifying selection (**Figure 7, Table S7**). For all patients, the same pattern of selection was found for blood and stool isolates, diversifying selection for patient A, neutral selection for patient B and purifying selection for patient C. The dN/dS estimation for patient B isolates, which are all mutators, is not indicative of the action of selection. However, the presence of mutators at high-frequency in the population (100 % in patient B) can be considered as a signature of sustained adaptation (57, 58). We also found that all isolates sampled in urine evolved under neutral selection (patients A and C). It should be noted that the number of structural variants (missense and synonymous) cannot be directly compared to dN/dS ratios. Structural variants were computed with reference to a single sequence whereas ratios were computed pairwise.

**Figure 7.**
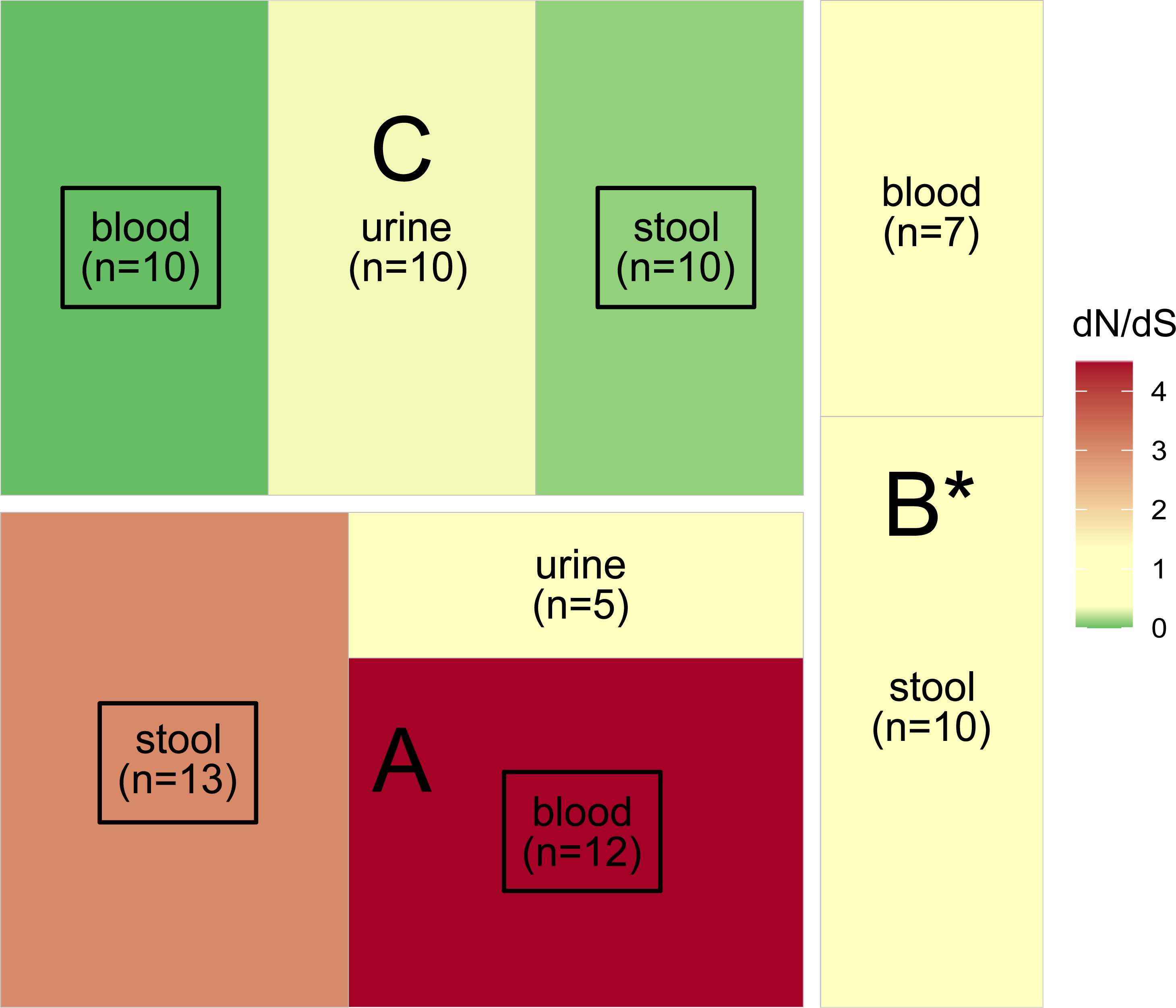
Action of selection on sequences (dN/dS) of *E. coli* isolates for patients A, B and C. Significant results are framed in black (see Table S7). All isolates of patient B are mutators (*). Neutrality (dN/dS not significantly different from 0) is indicated in pale yellow whereas purifying and diversifying selection are indicated in green and red, respectively.

## Discussion

*E. coli* is an opportunistic pathogen which may cross the intestinal barrier to reach extra-intestinal compartments causing infections. While evidence for purifying and neutral selection has been detected in commensal *E. coli* (10–12), extra-intestinal infection isolates have been shown to be under strong adaptive selection (13). Here we investigated both commensal and extra-intestinal isolates of leukemia patients undergoing chemotherapy. We observed the same strain in all compartments with a very limited number of SNPs, at the exception of the mutator strain. Interestingly, while each patient had a different selection signature, it was identical in both the bloodstream and the gut. In urine, in all cases, neutral selection was at play.

### Gut origin of urine and blood immunosuppressed patient isolates

For each patient, the isolates collected from the gut, urine, and blood exhibited highly similar sequences. They share the same phylogroup, ST, serotype, and *fimH* allele (**Table 1**), indicating that the strain most likely originated from the gut, as previously shown (34, 59). Moreover, with the exception of the mutator isolates, there were less than 15 SNPs between pairs of isolates of the same patient (**Figure 2**), indicating low genetic diversity, and the resistance and virulence profiles were stable (**Figures 5-6**). Interestingly, a previous study, on four patients, also showed a low diversity (ranging from 0 to 4 SNPs) when comparing urine isolates translocating to the blood (60). The phylogenetic distribution of isolates, together with the SNPs distribution among compartments, suggested the translocation of the gut population diversity to extra-intestinal compartments for all patients (**Figures 3-4**).

### Same adaptation forces at play in the gut and bloodstream of immunosuppressed patient

In non-immunosuppressed patients, purifying (or neutral) evolution of *E. coli* populations has been reported in the gut (12) but not in extra-intestinal compartments where bacteria DNA sequence are under diversifying selection with strong evidence for gene level convergence (13).

Here, we found various signatures of selection in gut isolates. In patient A, diversifying selection is at play, whereas purifying selection shaped the isolates of patient C. Differences in the microbiota perturbation of these patients undergoing a specific antibiotic therapy and chemotherapy could explain this discrepancy. However, the limited number of patients in this study prevents us from drawing such conclusions. The presence of mutators (all isolates of patent B) is indirect evidence for ongoing selection (57, 58). Some anticancer chemotherapy drugs enhance the bacterial mutagenesis, thus promoting the emergence of a mutator clone (61). Then, if the amount of beneficial mutations is large enough, a mutator clone can reach a high frequency.

Strikingly, the same adaptive process was at play in the bloodstream and in the gut for each patient. Moreover, unlike in extra-intestinal infections of non-immunosuppressed patients (13), no convergence among patients was detected. Finally, isolates were almost identical with a very low number of SNPs differentiating them (five on average) unspecific to the compartment. Whereas, in non-immunosuppressed patients, niche adaptation was evidenced by specific mutations associated with isolates from the gut or the bladder (28). These results suggest that the intestinal barrier does not act as a barrier anymore and that the adaptive constraints were homogenized among both compartments. A weakened immune system and a porous intestinal barrier, due to chemotherapy and antibiotic treatments, is the most obvious explanation for this phenomenon.

### Neutral selection in the bladder of immunosuppressed patient

For both patients with urine samples, isolates evolved neutrally. Fluctuating environment or a small population size could explain the weak strength of selection observed in the commensal habitat of *E. coli*. Indeed, bacteria in the bladder face continuous adaptive challenges with fluctuations in exposure to the host immune system, to antibiotic treatments, to nutrients and to a diverse microbial community in the host due to an alternation between storage and voiding phases (62).

### Limitations and future directions

Our work obviously presents a major limitation as we evaluated a very limited number of patients, which prevents us from drawing general conclusions. Collecting stool before antibiotic treatment is indeed very difficult to achieve in clinical practices. Nevertheless, the evaluation of larger cohorts with a greater number of samples is necessary to associate evolution patterns and the environment (patient follow-up and patient-dependent factors) and to decipher the factors linked to the selection patterns observed here.

## Conclusion

Our results highlight the complexity of the pathophysiology of *E. coli* BSIs. Without an efficient immune system and strong intestinal walls, these bacteria can easily cross the intestinal barrier and colonize extra-intestinal compartments. Here, we showed that these particular conditions homogenized the adaptive conditions in the gut and the bloodstream, further reinforcing the ease of large bacterial population translocation between them.

## Data availability

The data generated in this study have been submitted to the NCBI BioProject database under the accession number PRJEB69525 [https://www.ncbi.nlm.nih.gov/bioproject/?term=PRJEB69525].

## Code availability

The R scripts used to analyze the data and produce the figures are available on GitHub (https://github.com/j-marin/hemato_AVC) under a permissive licence (MIT).

## Acknowledgments

We are grateful to the INRAE MIGALE bioinformatics facility (MIGALE, INRAE, 2020. Migale bioinformatics Facility, doi: 10.15454/1.5572390655343293E12) for providing computing resources. We would also like to thank the Molecular Biology and Data Processing Platform of the Sorbonne Paris Nord University. This study was partly financed by two grants: one awarded by the research commission of the Université Sorbonne Paris Nord (BQR) and the other by the IFRB, UFR SMBH, Université Sorbonne Paris Nord.

